# AUTOENCODIX: powering a hub for biological representation learning models

**DOI:** 10.64898/2026.11.26.769509

**Authors:** Maximilian Joas, Kristina Müller, Alicia Diem, Luca Thale-Bombien, Christian Bäuchl, Aaron Klein, Jan Ewald

## Abstract

Autoencoders are a foundational architecture for biological representation learning and empower methods for denoising, multi-modal integration, explainable embeddings, and synthetic data generation. But, new architectures are scattered across isolated repositories, which hampers benchmarking, the reuse of pretrained models, and the transfer to new data modalities. Here, we present the next generation of AUTOENCODIX, a Python package and model hub for biological representation learning. Its modular, standardized API unifies preprocessing, training, hyperparameter optimization, evaluation, generation, and explanation across a growing set of architectures, now including disentangled, masked, semi-supervised, and 2D/3D image autoencoders. To demonstrate its use at scale, we tuned, trained, and benchmarked 15 large autoencoders on a balanced corpus of 3.7 million human cells from CELLxGENE, including nine ontology-based variational autoencoders whose latent dimensions represent core cellular functions and were defined by three large language models (LLMs). The biological-informed embeddings performed on par with classical Principal Component Analysis in downstream classification while being explainable by design, and in synthetic data generation they produced more biologically plausible error profiles than conventional variational autoencoders. Finally, AUTOENCODIX offers a new and unique combination of three pillars of explainability: by-design with biologically-informed latent dimensions, post-hoc gene-to-latent attribution scoring, and LLM-based knowledge integration for verbal description and hypothesis generation. AUTOENCODIX is open-source, open for community extensions and all pretrained models are available via Hugging Face for application on other single-cell gene expression data.

## INTRODUCTION

The rapid accumulation of high-throughput biomedical data has transformed the life sciences and created a need for computational methods that extract meaningful biological signals from high-dimensional data ^1^. Representation learning has become a cornerstone of this shift ^2^. Autoencoders (AEs) first gained traction as deep-learning-based alternatives for dimensionality reduction ^3^, but their value in biomedical research extends beyond replacing linear methods like Principal Component Analysis (PCA) or manifold learning techniques like Uniform Manifold Approximation and Projection (UMAP). AEs serve as a foundational architecture for a wide range of specialized methods that address central challenges in computational biology ^4^.

Modern AE architectures have proven valuable in several functional domains: (1) data imputation and denoising to handle the sparsity and technical noise of single-cell and spatial omics data ^5,6^; (2) multi-modal integration and modality translation to find a shared biological latent space across omics layers ^7,8^; (3) knowledge incorporation and explainable embeddings that map inputs into biologically constrained latent spaces reflecting known cellular pathways and networks ^9,10^; (4) generation of synthetic data ^11^ that enables federated, privacy-preserving machine learning (ML) ^12^; and (5) batch effect correction, cross-species integration ^13^, and perturbation modeling that separate technical covariates from biological variation and predict cellular responses to *in silico* genetic or chemical perturbations ^14^. Together, these capabilities support biological discovery, biomarker identification, disease prognosis, and the use of sensitive data through federated workflows.

Despite this success, the field lacks a unified software framework and a central model hub for biological representation learning. New AE architectures and methods are typically released in individual repositories with little standardization, generalization, or inter-operability. This hinders rigorous benchmarking, the exchange of pretrained models, and the adaptation of existing architectures to new data modalities ^15^. The first iteration of AUTOENCODIX partially addressed these challenges with an end-to-end pipeline for a curated set of AE architectures ^16^. It enabled robust benchmarking, systematic hyperparameter sensitivity studies, and user-friendly application interfaces. As a Makefile-based, monolithic pipeline of Python scripts, however, it lacked the flexibility, interoperability, and extensibility of a coherent Python package. Here, we present the next generation of AUTOENCODIX, an ecosystem and hub for biological representation learning, with three main contributions:

1. *Foundation: a standardized, modular Python package*. We re-implemented AUTOENCODIX as a Python package with a consistent API across all pipeline steps, inspired by standard frameworks like scikit-learn ^17^. Its decoupled, developer-centric codebase separates core components such as loss functions and network topologies from training routines and visualization. This allowed us to add new architectures and features, extend modality support to biomedical images, and support storing, sharing, and loading pretrained models.
2. *Application: large pretrained models*. We show how AUTOENCODIX can tune, train, and benchmark autoencoders on atlas-scale single-cell data, and we release 15 large pretrained models for scRNA-seq data, evaluated for embedding quality and synthetic data generation.
3. *Explainability: three complementary pillars of explainable AI (xAI) in one framework*. AUTOENCODIX combines a) by-design explainability via ontology-based autoencoders ^10,18^ that map latent dimensions to known biological pathways, b) post-hoc attribution via the Captum library ^19^ to identify the genes that drive specific latent dimensions, and c) LLM-based knowledge integration that turns these attributions into verbal, literature-backed descriptions of latent spaces to support hypothesis generation and biomarker discovery.

## RESULTS

### Feature overview and supported model architectures

To address the growing complexity and multi-modal nature of biological data, we turned AUTOENCODIX into a comprehensive Python package with an overhauled, modular, and flexible codebase designed for cross-platform usability and interoperability. It integrates with standard data structures and toolkits from scverse ^20^, PyTorch ^21^, and scikit-learn ^17^.

We expanded our autoencoder ‘zoo’ (Fig. 1A) to follow emerging trends and to support more data modalities, in particular images. Disentanglix is a variational autoencoder (VAE) that enforces control over the disentanglement of latent dimensions ^22,23^ and thereby helps to isolate distinct biological or technical factors of variation. Maskix implements a masked autoencoder for self-supervised learning, enabling data imputation and denoising ^24^. Since disease-related signals are often weaker than cell type or tissue variance, Supervisix extends the VAE in a semi-supervised manner with additional loss terms that control class separation and class cohesion, as introduced by Das *et al*. for speech emotion recognition ^25^. Finally, Imagix supports representation learning on 2D images and 3D volumes, such as medical images (e.g., computed tomography, CT, or magnetic resonance imaging, MRI) and spatial transcriptomics imaging. Its convolutional VAE learns compact representations that preserve spatial structure, as previously shown for cellular imaging ^7^, and the 3D extension preserves spatial context across the slices of volumetric images ^26^.

**Figure 1:**
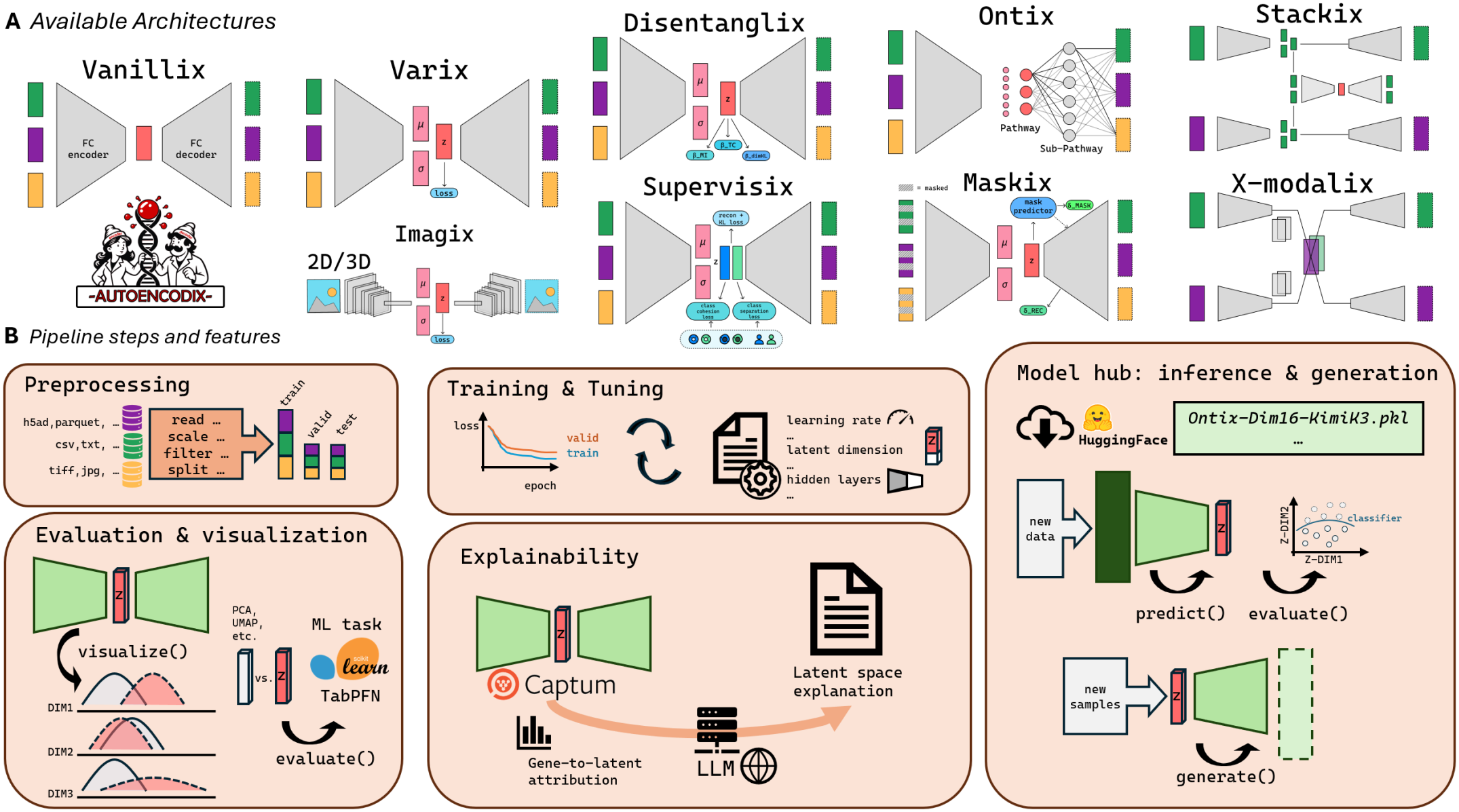
Overview AUTOENCODIX. **(A)** Sketches and list of available autoencoder architectures. **(B)** Key functionalities and pipeline steps covered by the framework.

AUTOENCODIX offers two new pipeline steps (Fig. 1B) that connect representation learning with downstream utility and inter-pretability. First, `generate()` simplifies synthetic data generation: users sample from the latent space and decode these samples into synthetic data of complex biological modalities. Second, `explain()` addresses the “black box” nature of deep learning embeddings with a dual-layer approach. It computes post-hoc feature-to-latent attribution scores via the Captum library and couples them with LLM-based integration of external knowledge, which verbalizes and contextualizes the biological mechanisms behind the attributions. We also extended existing functionalities. Data preprocessing now includes optimized support for single-cell data. Training can be coupled with any hyperparameter optimization (HPO) tool such as Syne Tune ^27^ or Optuna ^28^. Downstream evaluation of embeddings accepts any scikit-learn-like model, so that new tabular foundation models such as TabPFN ^29^ can be used directly on the embeddings.

AUTOENCODIX is designed for community-driven development and is ready for AI-agent-driven *autoresearch*^1^. The codebase follows the SOLID principles ^30^ (single responsibility, open/closed, Liskov substitution, interface segregation, dependency inversion), which promote modular and flexible software. The decoupling of components minimizes code duplication and speeds up the implementation of new architectures through module reuse. For example, we implemented Disentanglix simply by inheriting the pipeline of the baseline Varix model and supplying a modified loss function. Finally, AUTOENCODIX provides a standardized way to exchange autoencoder models in biological representation learning. Pipelines and network weights are self-contained objects, so researchers can share pretrained models tailored to specific biological contexts or data modalities. In the following, we showcase this by developing, benchmarking, and xAI with large pretrained ontology-based VAEs for single-cell RNA sequencing (scRNA-seq) data.

### Developing and benchmarking large pretrained autoencoders for scRNA-seq data

#### A balanced training corpus and LLM-defined ontologies

Like most large and foundation models for scRNA-seq data ^31–33^, we derived our training set from the CELLxGENE corpus ^34^. Instead of balancing at batch level during training, we defined and retrieved 4,158 chunks of unique combinations of cell type, tissue, sex, and disease state of human cells. We randomly downsampled each chunk to at most 1,000 cells, which resulted in a balanced corpus of about 3.7 million cells (Fig. 2A). We used this corpus to tune, train, and evaluate large AE models, focusing on ontology-based VAEs (Ontix) that provide explainable embeddings of high-dimensional scRNA-seq data. To this end, we used LLMs to define 16, 32, or 48 biological dimensions that reflect core cellular functions or disease-related processes and serve as latent dimensions. We further asked the LLMs to link these high-level terms to existing terms from the ontology databases KEGG, Reactome, or Gene Ontology (see Methods and the prompt in Supplementary Figure 3). The resulting two-layer ontology defines the sparse, biologically informed decoder and the gene input space.

**Figure 2:**
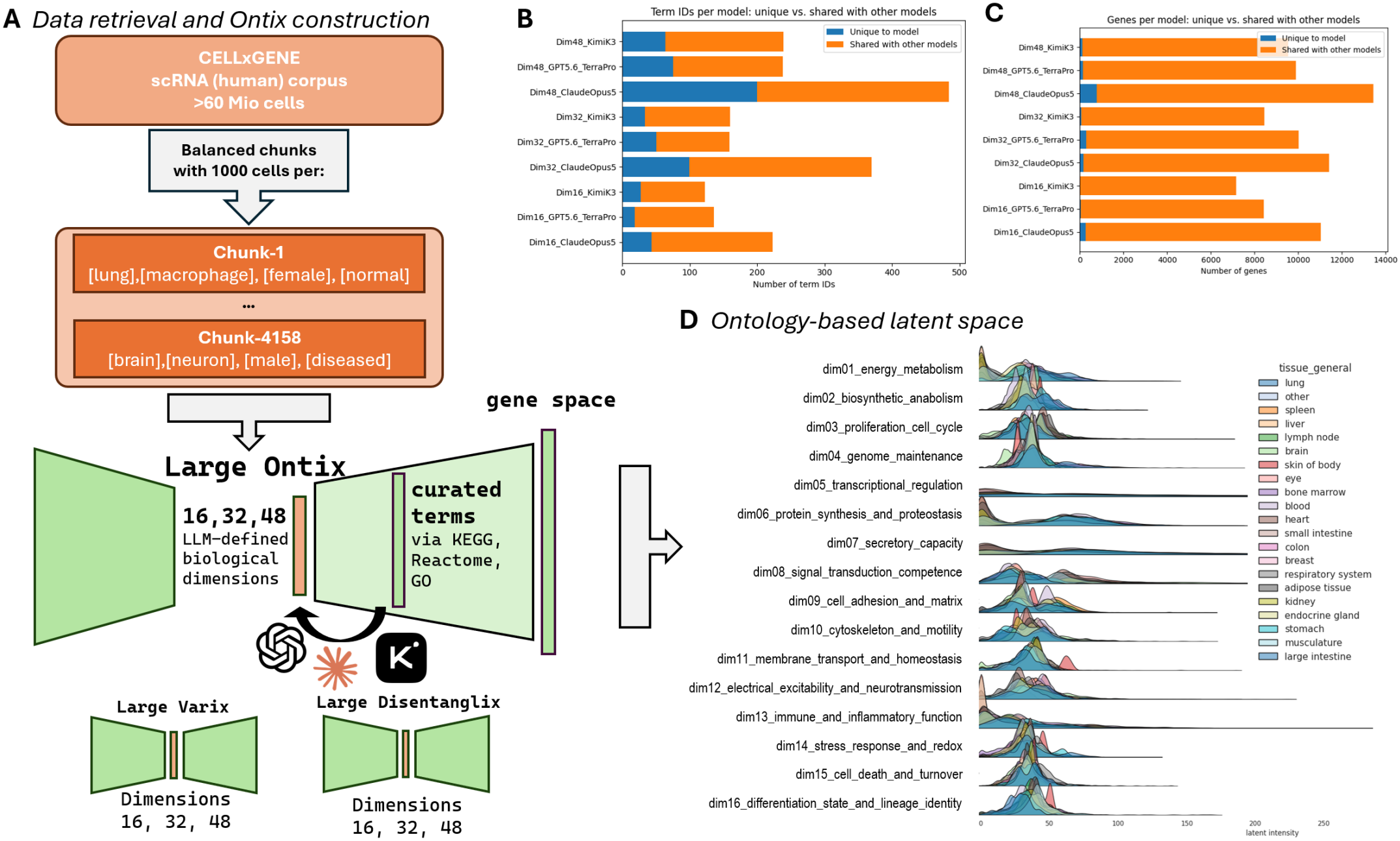
Large pretrained models. **(A)** Data retrieval and model construction workflow. **(B)** Number of ontology terms per Ontix model and **(C)** number of genes per ontology available for model building. **(D)** Example of a final latent space representation as ridgeline plot of latent intensities across tissues for model Dim16 Claude Opus 5.

The three LLMs (Claude Opus 5, GPT-5.6 TerraPro, and Kimi K3) shared the majority of ontology terms linked to their latent dimensions, with the 48-dimensional Claude Opus 5 ontology containing the most unique terms (Fig. 2B). The largest variation lies in the gene input space: Claude-derived ontologies cover a considerably larger portion of the genome, whereas Kimi K3 ontologies cover the smallest (Fig. 2C).

#### Tuning and training of autoencoders on large-scale data

We tuned and trained nine large Ontix models by combining three LLMs with latent dimensions of 16, 32, or 48. We tuned the hyperparameters of each model individually to fully exploit the architecture, which is composed of five fully connected encoder layers and two sparsely connected decoder layers. Using Syne Tune with downstream task performance as the optimization objective, similar to our previous benchmarks ^16^, we varied the loss-term weight *β*, training parameters (learning rate, batch size, dropout rate, weight decay), and the encoding factor (see Hyperparameter optimization and model training for details).

HPO revealed several insights into parameter sensitivity. Performance converged rapidly after 50 to 100 HPO trials (Supplementary Figure 1A). Downstream task performance and reconstruction loss were negatively correlated, though not perfectly, with Spearman rank correlations between *−*0.4 and *−*0.7 (Supplementary Figure 1B). The learning rate was highly sensitive: higher values drove faster optimization and higher downstream utility at the final epoch, but beyond a critical threshold, training became unstable and utility collapsed (Fig. 3A). Both the dropout rate and *β* were consistently optimized toward low values, i.e., toward weaker regularization of encoder and latent space. In contrast, weight decay had a negligible impact on the objective, likely because the AdamW optimizer decouples weight decay from the learning rate ^35^. The optimal encoding factor, i.e., the degree of shrinkage within the encoder, ranged from about 1.0 to 1.7. Overall, the optimal configurations (Table 1) were largely consistent across ontologies. Only Dim16_ClaudeOpus5 was an exception that combined a high dropout rate with a wide encoder (encoding factor close to one).

**Table 1.** Architecture statistics and hyperparameter configuration of the trained models. Parameter counts are given in millions (M); “non-zero” denotes the parameters remaining after sparsification and weights with absolute values *≥*0.01. Hidden-layer sizes are listed from input to latent space (encoder) and from latent space to output (decoder). Decoder sparsity is given per decoder layer.

| Model | Params. total (M) | Params. non-zero (M) | Input dim. | Encoder hidden | Latent dim. | Decoder hidden | Decoder sparsity |
|---|---|---|---|---|---|---|---|
| Dim16_ClaudeOpus5 | 373.28 | 365.68 | 9006 | 8845, 8687, 8531, 8378, 8228 | 16 | 207 | 0.9369, 0.9907 |
| Dim32_ClaudeOpus5 | 151.67 | 138.23 | 10331 | 7339, 5213, 3703, 2630, 1868 | 32 | 332 | 0.9687, 0.9935 |
| Dim48_ClaudeOpus5 | 520.27 | 370.58 | 12499 | 11420, 10434, 9534, 8711, 7959 | 48 | 446 | 0.9791, 0.9949 |
| Dim16_KimiK3 | 27.39 | 26.60 | 5401 | 3216, 1915, 1140, 679, 404 | 16 | 109 | 0.9375, 0.984 |
| Dim32_KimiK3 | 88.01 | 86.44 | 7396 | 5510, 4105, 3058, 2278, 1697 | 32 | 147 | 0.9688, 0.9864 |
| Dim48_KimiK3 | 384.72 | 300.49 | 9462 | 9157, 8862, 8576, 8300, 8032 | 48 | 228 | 0.9791, 0.9898 |
| Dim16_GPT5.6_TerraPro | 86.95 | 85.14 | 8357 | 5666, 3841, 2604, 1765, 1196 | 16 | 124 | 0.9375, 0.9862 |
| Dim32_GPT5.6_TerraPro | 105.14 | 101.51 | 9749 | 6287, 4054, 2614, 1685, 1086 | 32 | 143 | 0.9685, 0.989 |
| Dim48_GPT5.6_TerraPro | 101.90 | 96.41 | 9827 | 6182, 3889, 2446, 1538, 967 | 48 | 223 | 0.979, 0.9916 |
| Dim16_Varix | 335.43 | 303.21 | 16375 | 8187, 4093 | 16 | 4093, 8187 | 0, 0, 0 |
| Dim32_Varix | 335.63 | 308.39 | 16375 | 8187, 4093 | 32 | 4093, 8187 | 0, 0, 0 |
| Dim48_Varix | 335.82 | 309.78 | 16375 | 8187, 4093 | 48 | 4093, 8187 | 0, 0, 0 |
| Dim16_Disentanglix | 335.43 | 304.41 | 16375 | 8187, 4093 | 16 | 4093, 8187 | 0, 0, 0 |
| Dim32_Disentanglix | 335.63 | 311.12 | 16375 | 8187, 4093 | 32 | 4093, 8187 | 0, 0, 0 |
| Dim48_Disentanglix | 335.82 | 310.87 | 16375 | 8187, 4093 | 48 | 4093, 8187 | 0, 0, 0 |
| Model | Learning rate | Weight decay | Epochs | Batch size | Dropout rate | Encoding factor | β |
| Dim16_ClaudeOpus5 | 2.50×10<sup>-3</sup> | 1.88×10<sup>-4</sup> | 500 | 1659 | 0.554 | 1.02 | 1.66×10<sup>-5</sup> |
| Dim32_ClaudeOpus5 | 3.60×10<sup>-3</sup> | 2.04×10<sup>-3</sup> | 500 | 3339 | 0.028 | 1.41 | 7.06×10<sup>-5</sup> |
| Dim48_ClaudeOpus5 | 1.81×10<sup>-3</sup> | 3.64×10<sup>-2</sup> | 500 | 2276 | 0.153 | 1.09 | 1.13×10<sup>-4</sup> |
| Dim16_KimiK3 | 3.88×10<sup>-3</sup> | 1.90×10<sup>-5</sup> | 500 | 1235 | 0.096 | 1.68 | 4.94×10<sup>-5</sup> |
| Dim32_KimiK3 | 4.42×10<sup>-3</sup> | 1.64×10<sup>-5</sup> | 500 | 1181 | 0.074 | 1.34 | 3.35×10<sup>-5</sup> |
| Dim48_KimiK3 | 2.26×10<sup>-3</sup> | 1.90×10<sup>-3</sup> | 500 | 958 | 0.182 | 1.03 | 1.27×10<sup>-5</sup> |
| Dim16_GPT5.6_TerraPro | 6.15×10<sup>-3</sup> | 1.10×10<sup>-5</sup> | 500 | 3097 | 0.055 | 1.47 | 2.20×10<sup>-5</sup> |
| Dim32_GPT5.6_TerraPro | 3.84×10<sup>-3</sup> | 3.36×10<sup>-5</sup> | 500 | 2348 | 0.030 | 1.55 | 8.49×10<sup>-5</sup> |
| Dim48_GPT5.6_TerraPro | 2.22×10<sup>-3</sup> | 1.07×10<sup>-3</sup> | 500 | 1332 | 0.065 | 1.59 | 2.07×10<sup>-5</sup> |
| Dim16_Varix | 1.00×10<sup>-3</sup> | 1.00×10<sup>-3</sup> | 500 | 2048 | 0.100 | 2.00 | 1.00×10<sup>-4</sup> |
| Dim32_Varix | 1.00×10<sup>-3</sup> | 1.00×10<sup>-3</sup> | 500 | 2048 | 0.100 | 2.00 | 1.00×10<sup>-4</sup> |
| Dim48_Varix | 1.00×10<sup>-3</sup> | 1.00×10<sup>-3</sup> | 500 | 2048 | 0.100 | 2.00 | 1.00×10<sup>-4</sup> |
| Dim16_Disentanglix | 1.00×10<sup>-3</sup> | 1.00×10<sup>-3</sup> | 500 | 2048 | 0.100 | 2.00 | 0.1 |
| Dim32_Disentanglix | 1.00×10<sup>-3</sup> | 1.00×10<sup>-3</sup> | 500 | 2048 | 0.100 | 2.00 | 0.1 |
| Dim48_Disentanglix | 1.00×10<sup>-3</sup> | 1.00×10<sup>-3</sup> | 500 | 2048 | 0.100 | 2.00 | 0.1 |

**Figure 3:**
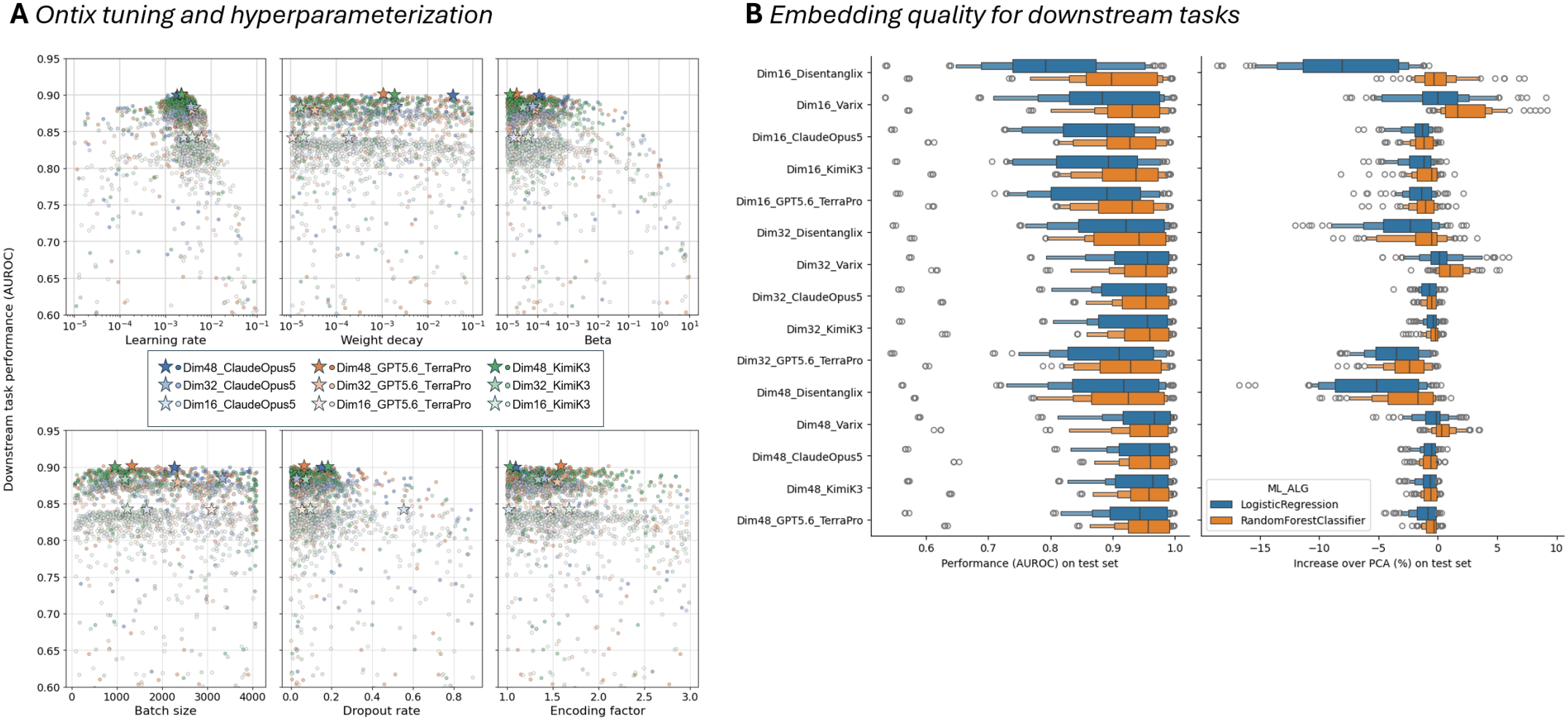
Tuning and evaluation results. **(A)** Scatter plot of individual tuning runs (points) for each hyperparameter value (x-axis) vs. average downstream task performance of embeddings (y-axis). Stars represent best configurations per model. **(B)** Downstream task performance of embeddings generated and evaluated with final trained models with logistic regression and RandomForest classifiers. Performance is displayed as AUROC (left) and as increase over PCA embeddings with similar dimensionality (right).

Based on the tuning results, we trained the final Ontix models with their optimal configurations for twice as many epochs. For the baseline Varix and Disentanglix models, we set the hyperparameters manually, guided by the optimal Ontix configurations. We trained all final models on a single NVIDIA H100 GPU with a low VRAM footprint (<12 GB) by converting sparse single-cell data to dense tensors only at batch level.

#### Embedding quality

A typical use case for pretrained autoencoders is to generate explainable, low-dimensional embeddings of high-dimensional, sparse scRNA-seq data (Fig. 2D). Such embeddings are used for visualization, clustering, and downstream classification, such as the prediction of cell type or disease state. We evaluated the latter on a defined set of tasks based on available cell annotations (Supplementary Table 1). For each task, we compared a linear model (logistic regression) and a simple non-linear model (random forest) trained on our embeddings and on PCA embeddings of the same dimensionality (Fig. 3B; results per task in Supplementary Figure 2). As expected, higher latent dimensionality generally improved downstream performance. Random forests tended to perform slightly better than logistic regression, though not in all settings. Overall, performance was comparable across architectures. Conventional VAEs (Varix) showed a slight advantage, most notably in the 16-dimensional latent space, where their improvement over PCA was strongest. The Ontix variants showed no consistent differences among each other and performed roughly on par with PCA. Thus, Ontix embeddings often do not achieve the best downstream performance, likely due to their restricted, linear decoder, but they offer built-in explainability in return.

#### Synthetic data generation with pretrained models

Beyond encoding, we assessed the models for synthetic data generation and compared VAEs with and without biologically informed latent dimensions. Our benchmark follows a simple rationale: a cell annotation tool such as CellTypist ^36^ should assign the same cell type to a cell before and after it is encoded (`predict()`) and decoded (`generate()`), as outlined in Figure 4A. For a broad utility assessment we used several CellTypist models specialized for different tissues or functional contexts, including immune cells.

**Figure 4:**
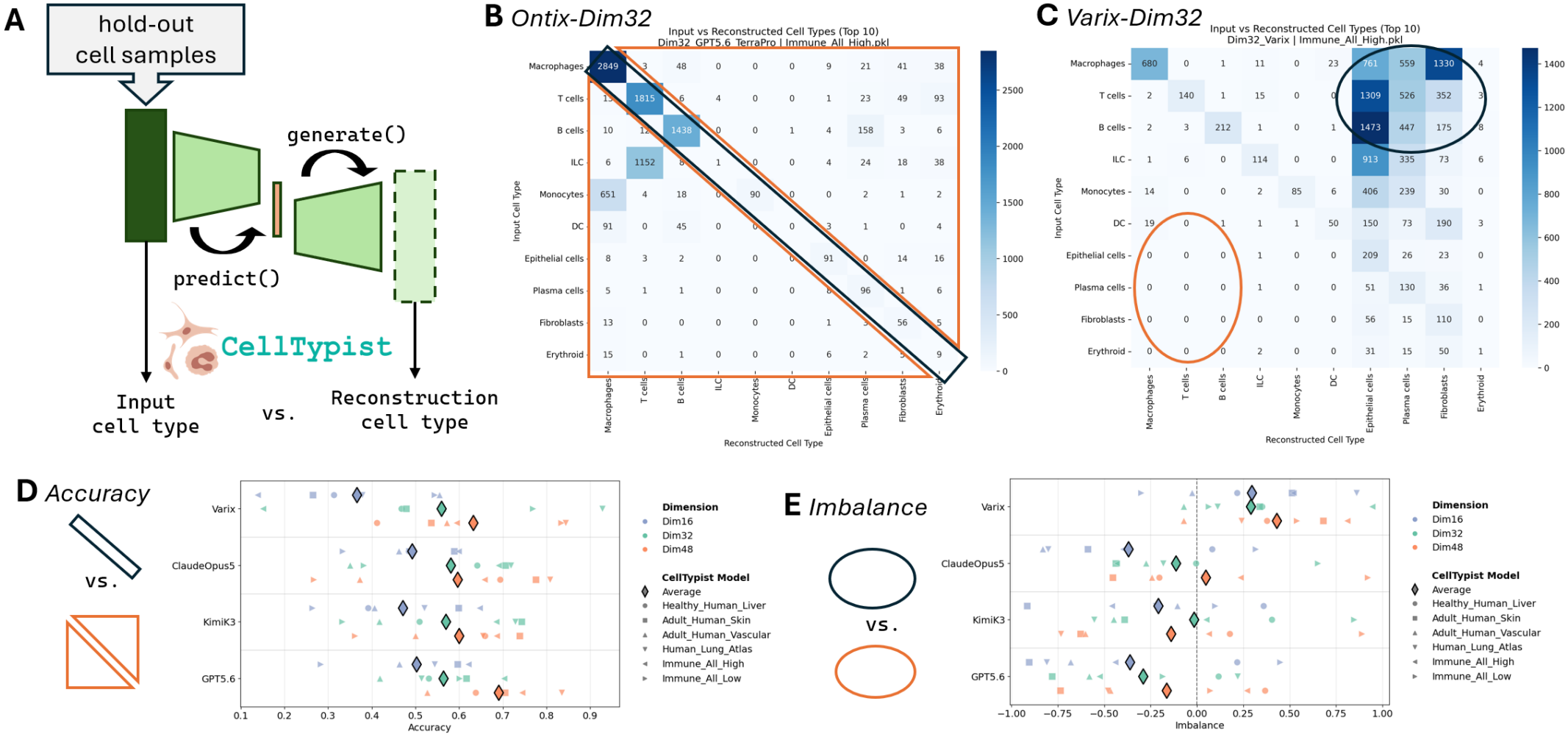
Benchmarking pretrained models for synthetic data generation. **(A)** Rationale behind the benchmark using cell type annotation with CellTypist. **(B)** and **(C)** Exemplary confusion matrices comparing cell type annotation before and after application of pretrained models to hold-out cells. **(D)** Accuracy and **(E)** imbalance of confusion matrices across all pretrained models and selected CellTypist models.

On average, conventional VAEs achieved comparable or lower raw accuracy in cell type reconstruction, and accuracy mostly increased with the number of latent dimensions. Among the Ontix models, Dim48 GPT-5.6 TerraPro showed the highest average accuracy, with substantial variation across tissues and CellTypist models but only small differences between LLMs. The confusion matrices revealed pronounced differences in the type of errors made by Ontix and Varix models. Ontix models tended to produce biologically plausible confusions, for instance shifting the gene expression profiles of monocytes toward those of macrophages (Fig. 4B). Because macrophages were not shifted toward monocytes to the same extent, the confusion matrix was imbalanced. In this example, the imbalance was predominantly negative, reflecting a collapse toward fewer cell types after reconstruction. In contrast, Varix models showed a strongly positive imbalance for high-level immune cell reconstruction (Fig. 4C and E) and shifted the profiles of macrophages toward several distinct, biologically unrelated cell types, such as epithelial cells or fibroblasts. Although this phenomenon warrants further investigation, it suggests that Ontix models produce a more biologically plausible error profile in synthetic data generation than conventional VAEs and that raw accuracy alone can be misleading.

### Explainability: combining by-design, post-hoc, and LLM-based xAI

Combining large Ontix models with the xAI functionalities of AUTOENCODIX uniquely integrates three complementary levels of explainability. First, the latent dimensions of Ontix models are directly associated with defined biological functions. Figure 5A illustrates this for normal lung cells and lung adenocarcinoma cells, which differ markedly along latent dimensions related to energy metabolism and adaptive immunity, consistent with the expected contrast between healthy and cancer cells. Second, post-hoc attribution scores reveal the contribution of features to specific latent dimensions and thereby uncover candidate biomarker gene sets. Third, LLMs integrate and verbalize biological knowledge about the identified gene sets. Figure 5B exemplifies this: we used Captum with the Integrated Gradients algorithm to identify the genes most influential in distinguishing healthy from cancer cells along the latent dimension capturing energy metabolism. We then passed the 15 most influential genes in a structured prompt to an LLM API of the user’s choice (here Gemini 3.8 Flash via OpenRouter), which generated a summary and biologically motivated hypotheses explaining the group differences. Notably, LLMs do not receive the latent dimension name and the summarized biological themes describing energy metabolism were purely derived from the most influential gene list, which shows its potential for hypothesis generation.

**Figure 5:**
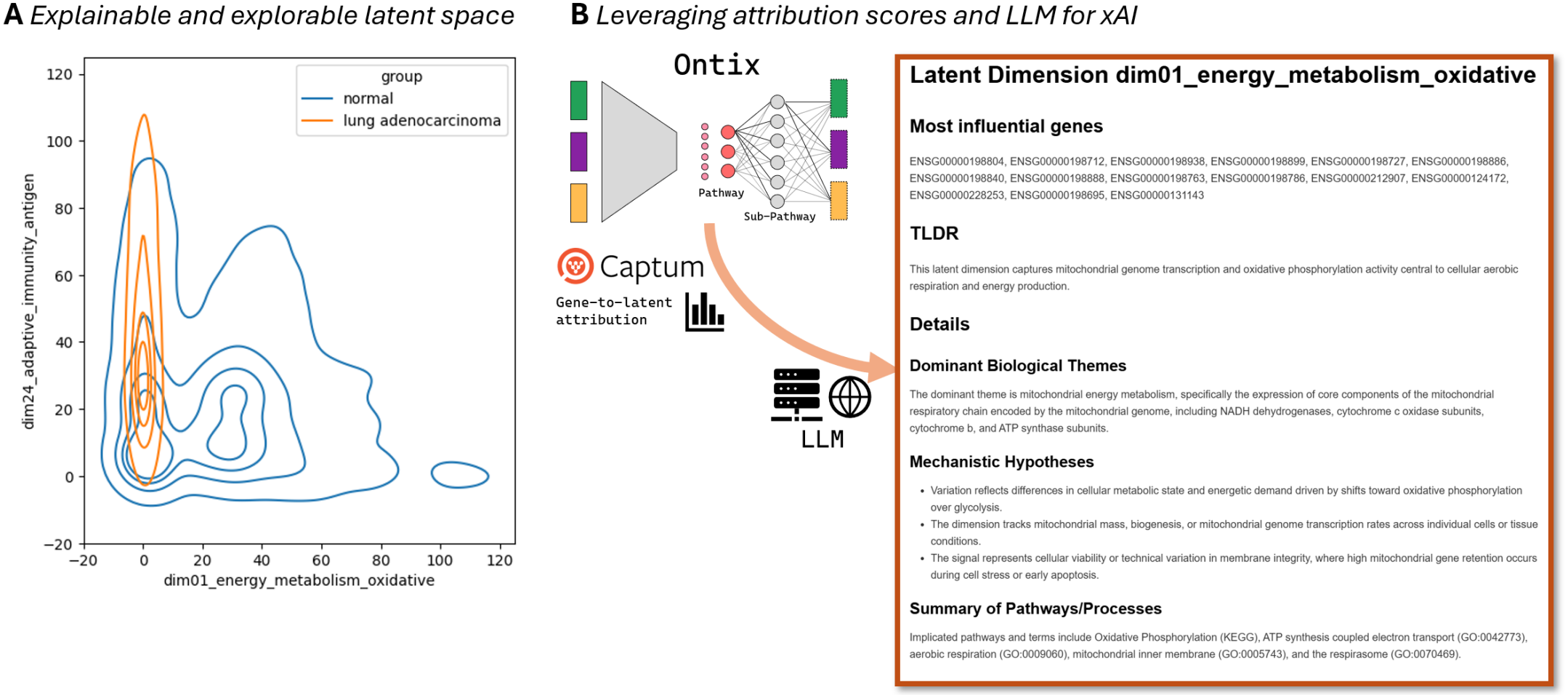
xAI with AUTOENCODIX. **(A)** Latent intensity distribution across two directly explainable latent dimensions using the Ontix model Dim32 Claude Opus 5 for healthy and cancer cells. **(B)** Feature-to-latent attribution scoring per latent dimension is integrated with LLMs (e.g., Gemini 3.8 Flash via OpenRouter) to generate a verbal description of the latent space as markdown file.

## FUTURE DIRECTIONS

We see several directions to extend AUTOENCODIX in terms of architectures, automated model configuration, and data applicability. First, we plan to support learning paradigms beyond reconstruction, notably contrastive learning. Contrastive approaches align similar samples and separate dissimilar ones in latent space and are particularly effective for modeling perturbation responses and condition-specific effects in single-cell data. A key example is the Compositional Perturbation Autoencoder (CPA), which combines an AE backbone with a compositional latent structure to disentangle drug, genetic, and covariate effects and to predict unseen perturbation combinations ^14^. Integrating contrastive and compositional objectives will extend AUTOENCODIX to perturbation modeling and causal-style representation learning.

Second, we aim to expand the pretrained model zoo across modalities (e.g., single-cell transcriptomics, proteomics, epigenomics) and tasks (e.g., cell type annotation, perturbation response modeling, biomarker discovery). This will lower the barrier for users working with non-standard data types and facilitate systematic benchmarking of ontology-informed versus standard architectures.

Third, we intend to use the AUTOENCODIX hub for meta-learning, building on BBOmix, an HPO benchmark comprising four architectures and seven omics modalities ^37^. As trained models, datasets, and performance metrics accumulate, the hub will enable learning relationships between dataset properties, architecture choices, and optimal hyperparameters. Such a meta-learning layer can warm-start hyperparameter searches and eventually enable zero-shot configuration based on dataset characteristics, moving toward parameter-free autoencoder training.

Fourth, we plan to extend AUTOENCODIX beyond standard autoencoders to broader self-supervised paradigms, particularly the Joint-Embedding Predictive Architecture (JEPA) ^38^. By predicting latent representations instead of reconstructing raw inputs, JEPA avoids the computational overhead and noise sensitivity of feature-level reconstruction, as shown for images (I-JEPA) ^39^ and videos (V-JEPA) ^40^. We also aim to incorporate expressive components like transformer self-attention blocks, inspired by single-cell foundation models such as scGPT ^31^ and Geneformer ^41^, to better capture complex gene–gene and gene–cell interactions.

Finally, we plan to broaden multi-modal support, for example to time-series data and to architectures that integrate imaging and omics data ^42^. This will enable integrated views of cellular state that combine molecular, structural, and temporal dynamics.

## MATERIALS AND METHODS

AUTOENCODIX is available as Python package via https://pypi.org/project/autoencodix/, its code is maintained at https://github.com/jan-forest/autoencodix_package/ and pretrained models are shared via https://huggingface.co/collections/autoencodix/acx-pretrained-models. Additional code to create data corpus, to train and to evaluate models is available in a dedicated repository for reproducibility: https://github.com/jan-forest/ACX_largemodel_reproducibility.

### Data and ontologies

#### Corpus creation

A human scRNASeq corpus was collected from the CZ CELLxGENE corpus using the CENSUS API at LTS version 2025-11-08. To avoid an over- and under-representation of cell types, tissues and other biases, we defined 4158 chunks as unique combination of available cell annotation parameters:

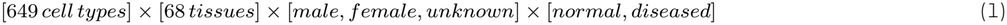

A balanced corpus was subsequently reached by first filtering out cells with less than 5% expressed genes and then random downsampling to 1000 cells per chunk and omitting chunks with less than 1000 cells. Each chunk is retrieved as AnnData file with annotation parameters and expression values are *log*1*p* transformed to avoid bias towards genes with very high expression when models are trained with Mean-Squared-Error (MSE) as reconstruction loss term. After preprocessing and joining AnnData files of all chunks, the total corpus of 3.7 million cells is randomly split into three splits with 5% for tuning, 90% for final training of models and 5% hold-out set to perform model evaluations.

#### Ontology generation with LLMs

We leveraged three state-of-the-art LLMs (in mid 2026) to create an ontology layer for the large pretrained Ontix-models. In comparison to rely solely on existing curated ontologies, this enables flexibility over the number of latent dimensions, the focus of ontology-based latent dimension and the advantage to combine multiple ontology databases. All LLMs (Claude Opus 5, GPT-5.6 TerraPro and Kimi K3 via OpenRouter) were asked using the same prompt (see Supplemental Figure 3) to specify a JSON-output containing 16, 32 or 48, distinct biological and cellular dimensions where each is assigned to 5-20 existing ontology-terms from sources KEGG, Reactome or Gene Ontology (without conceptual overlap). The second ontology layer is then completed by mapping ontology-terms from KEGG, Reactome and Gene Ontology to HUGO symbols and ENSEMBL gene identifiers using MyGene package. Resulting model and ontology statistics are provided in Table 1.

### Hyperparameter optimization and model training

#### Tuning

To optimize architectural and training hyperparameters as well as the loss term weight *β*, we used Syne Tune and the smaller tuning split (5% of total) for HPO by using 106,246 cells (60%) as training set and 63,747 as validation set (40%). In each run models are trained for 250 epochs using the loss function and then evaluated on a broad collection of downstream tasks leveraging both given annotation parameters like tissue source and 25 LLM-generated cell type classification tasks (see Supplemental Table 1). Hyperparameters are then optimized by maximization of the average across annotation tasks and cell type tasks. HPO is performed individually for each ontology and training and evaluating for a budget of 48h in parallel on 12x NVIDIA H100 resulting in 147-276 completed tuning trials.

#### Final model training

Final Ontix models are trained on a single NVIDIA H100 and on the training split containing 3,330,818 cells and with hyperparameters determined by tuning for an increased number of 500 epochs.

For comparison Varix and Disentanglix models were trained accordingly. For non-ontology-based models, hyperparameterization (see Table 1) was defined manually based on the observed optimal values of Ontix providing a computational efficient comparison.

### Evaluation and xAI methods

For all evaluation steps the hold-out set of 186,396 cells was used which was not part of tuning nor training of the models. Using the `predict()` pipeline step in AUTOENCODIX, latent space embeddings were calculated serving as foundation for latent space visualizations and evaluation on downstream machine learning task via pipeline step `evaluate()`.

#### Downstream task

Evaluation was performed similarly to hyperparameter tuning on a broad range of classification tasks based on annotation parameters like tissue and 25 LLM-generated cell type classification tasks (see Supplemental Table 1). The `evaluate()` is designed to handle any machine learning algorithm with scikit-learn-like API and we evaluated embeddings for simplicity with the linear classifier `LogisticRegression()` and the non-linear classifier `RandomForestClassifier()`. To avoid overfitting the RandomForest models are trained with only 100 trees, maximum depth of eight nodes and a minimum of four samples per leaf. Model performance was assessed via three-fold cross-validation and using PCA-based embeddings with similar dimensionality as reference method to autoencoders. For a fair comparison PCA is fit prior to the same data split used for tuning before applied to the hold-out split.

The downstream task performance is compared across dimensions by calculating z-score representing the mean (*avg*) and standard deviation (*sd*) normalized increase (positive) or decrease (negative) in comparison to PCA across three folds (*r ∈* [1, 3]) using the following equation:

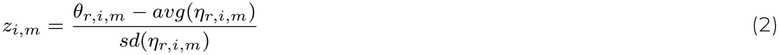

where *θ*_*r,i,m*_ is the performance of the embedding of the *m*-th autoencoder model for the *i*-th classification task as ROC AUC and *η*_*r,i,m*_ the corresponding PCA embedding performance.

#### Latent space xAI methods

In the AUTOENCODIX pipeline step `explain()` we combine *post-hoc* xAI attribution scoring to reveal feature-to-latent dimension contribution via Captum integration, with LLM-based enrichment with biomedical knowledge about the identified most influential features (genes) per latent dimension.

In the provided example Integrated Gradients is used as attribution scoring algorithm to determine the 15 most influential genes per dimension comparing lung adenocarcinoma cells (target) to non-diseased lung cells (baseline) on a subset of 400 randomly selected samples per group from the hold-out split. A LLM-based summary for each dimension and their 15 most influential genes is then generated using Google Gemini Flash 3.8 using our default prompt template which queries a structured JSON (see https://github.com/jan-forest/autoencodix_package/blob/main/src/autoencodix/utils/prompts.py). For inference we used OpenRouter.

#### Data generation evaluation

To check model capabilities generating synthetic data from user-defined priors of latent space, we leveraged CellTypist models as common cell type annotation tool. To assess model biases when generating data, we first calculate latent space embeddings of a subset of cells from the hold-out split matching the domain of the selected CellTypist model. For a maximum of randomly downsampled 100,000 cells, the gene expression profile is reconstructed via the pipeline step `generate()` by using the embedding as priors for the decoder. Finally, gene expression profiles of input and reconstruction are each annotated with the respective CellTypist model and compared to generate confusion matrices. Accuracy is then defined as the proportion of the diagonal to all entries in the confusion matrix. Since we noticed a bias of models and non-symmetric distribution of errors, we calculate an imbalance score by comparing the sum of the upper triangle off the diagonal with the lower one:

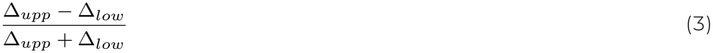

where zero shows a balanced confusion matrix and +1 respectively *−*1 maximally imbalanced cell type annotation errors. A positive imbalance score represents the fact that an input cell type class like macrophages is misclassified over many other classes (*expansion*), whereas a negative imbalance score represents that multiple input classes, like macrophages plus monocytes, are classified as single class (macrophages) on the reconstructed gene expression profile (*contraction*).

## DATA AVAILABILITY

All data used in this study are publicly available. The single-cell RNA-seq data were obtained from the CZ CELLxGENE Census ^34^ (LTS release 2025-11-08, *Homo sapiens*). The balanced corpus of 3.7 million cells derived from it, with its train, tune and hold-out splits, is available as AnnData files and as AUTOENCODIX data containers at https://huggingface.co/datasets/autoencodix/census-large. All 15 pretrained models (Ontix, Varix and Disentanglix with 16, 32 and 48 latent dimensions) are available, together with the LLM-generated ontologies, at https://huggingface.co/collections/autoencodix/acx-pretrained-models. Extended results are deposited at Zenodo (https://doi.org/10.5281/zenodo.24396083), including hyperparameter tuning results, hold-out latent embeddings, PCA baselines, downstream evaluation tables, CellTypist comparisons and plots. The CellTypist models used for the data generation benchmark are publicly available at https://www.celltypist.org/models.

## CODE AVAILABILITY

AUTOENCODIX is available as a Python package at https://pypi.org/project/autoencodix/ (version 0.2.4 was used to train models and generate results), and its source code is at https://github.com/jan-forest/autoencodix_package. Additional code to create the corpus, generate the ontologies, and tune, train and evaluate the models, and to reproduce all figures, is available at https://github.com/jan-forest/ACX_largemodel_reproducibility. That repository also contains the LLM prompts and outputs, the final ontologies, and the definitions of the classification tasks.

## ACKNOWLEDGMENTS

JE, MJ, AD and LTB acknowledge financial support by the Federal Ministry of Education and Research of Germany (BMFTR) and by Sächsisches Staatsministerium für Wissenschaft, Kultur und Tourismus in the programme Center of Excellence for AI-research „Center for Scalable Data Analytics and Artificial Intelligence Dresden/Leipzig”, project identification number: ScaDS.AI. KM acknowledges support by the BMFTR under the funding code 03ZU2111DC (SaxoCell NK-Alliance) as part of the Clusters4Future cluster SaxoCell. LTB acknowledges support by the BMFTR through a scholarship of DAAD project 57616814 (SECAI, School of Embedded and Composite AI) as part of the program Konrad Zuse Schools of Excellence in Artificial Intelligence. The authors gratefully acknowledge the computing time made available to them on the high-performance computer at the NHR Center of TU Dresden.

## AUTHOR CONTRIBUTIONS

MJ drafted the software and framework code and JE continued software development. AD, LTB, CB, KM contributed with features and extensions of software. JE and AK supervised developments. JE was mainly responsible for model training, experiment design and data analysis with support from all authors. JE wrote manuscript draft and all authors contributed in writing the final manuscript including visualizations and repository preparation.

## SUPPLEMENTARY INFORMATION

**Supplementary Figure 1:**
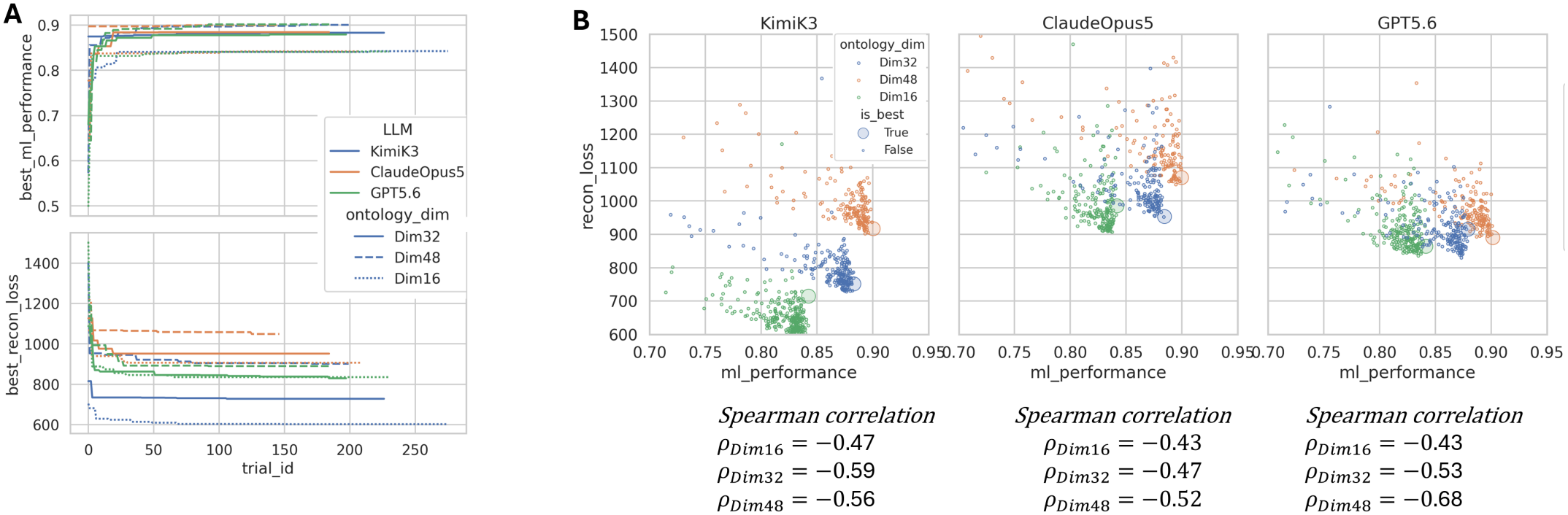
Details of tuning. **(A)** Evolution of embedding quality (downstream task performance) and reconstruction loss on valid set along tuning trials. **(B)** Correlation of embedding quality and reconstruction loss for each model.

**Supplementary Figure 2:**
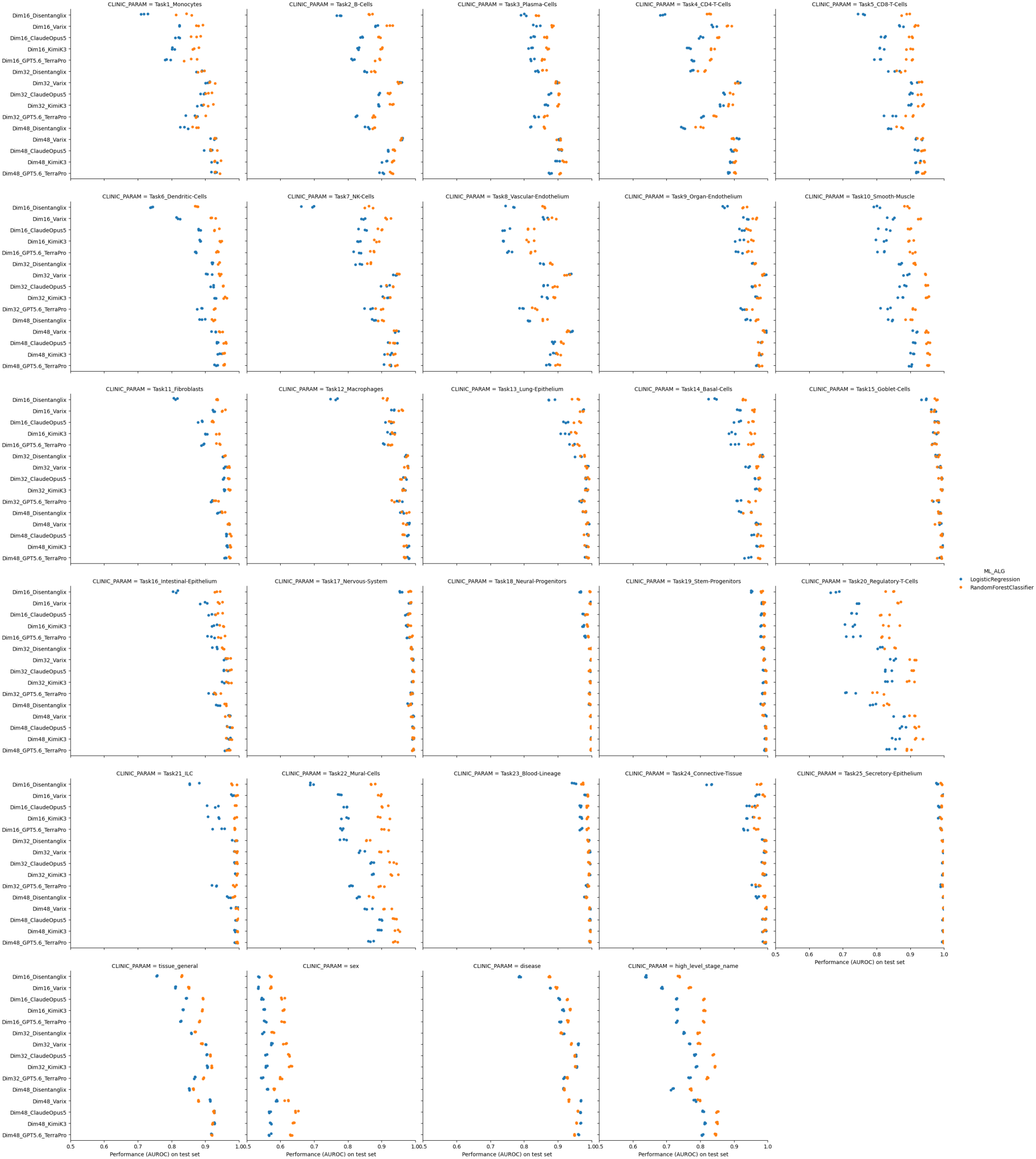
Detailed embedding quality. Embedding quality as downstream task performance shown individually for each task across all models on the hold-out test cell samples.

### Prompt template to generate ontologies with LLMs

**Supplementary Figure 3:**
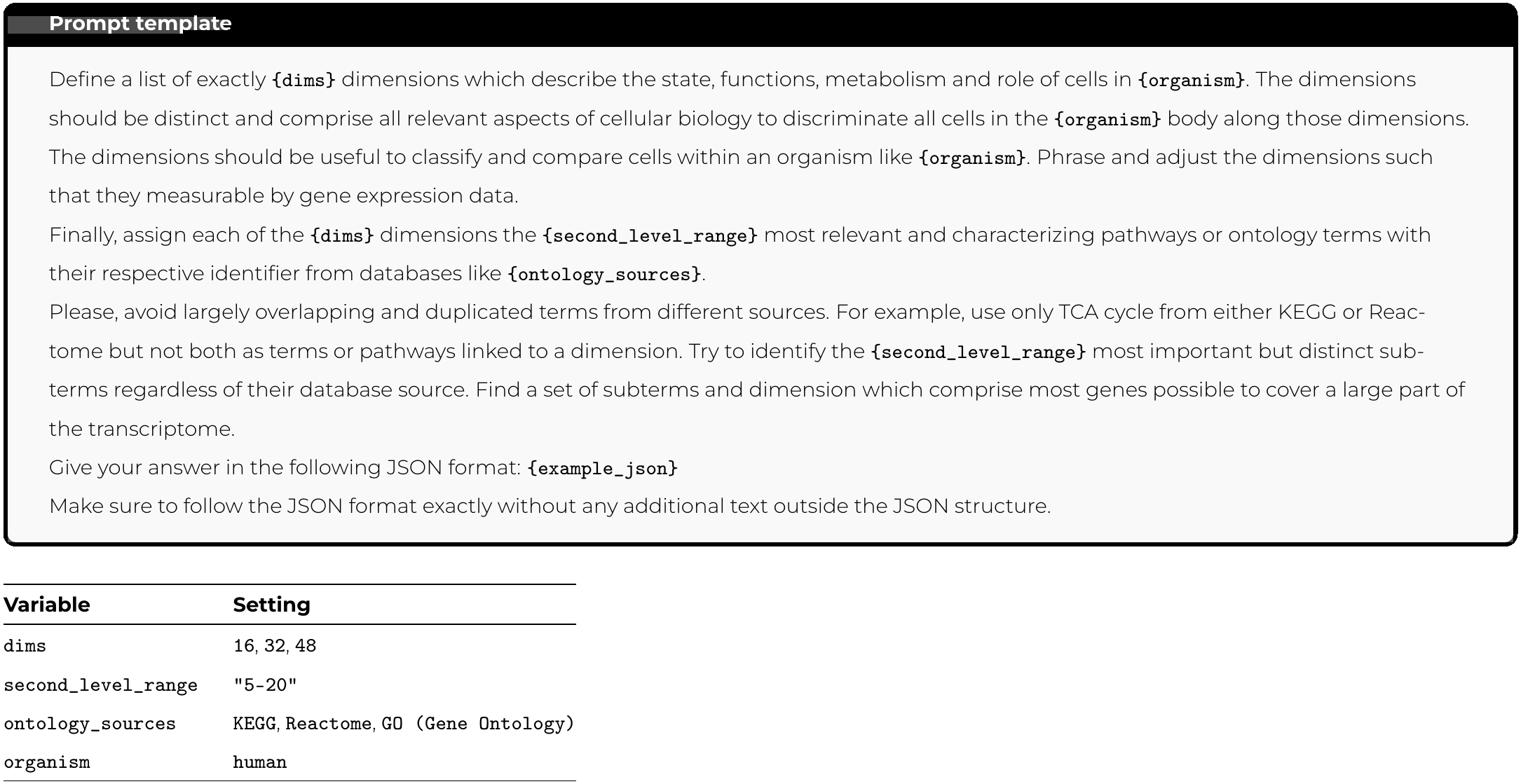
Prompt template and variable settings used to query the LLM. Monospaced terms in braces are placeholders filled programmatically. `example_json` is a schema example that defines the output format.

### Classification tasks

**Supplementary Table 1:** Cell-type classification tasks and metadata prediction tasks.

| **#** | **Task** | **Cell types** |
|---|---|---|
| 1 | Monocytes | classical monocyte; non-classical monocyte; CD14-positive monocyte; CD14-positive, CD16-positive monocyte |
| 2 | B cells | naive B cell; memory B cell; mature B cell; germinal center B cell |
| 3 | Plasma cells | IgA plasma cell; IgG plasma cell; plasmablast; plasma cell |
| 4 | CD4 T cells | naive thymus-derived CD4-positive, alpha-beta T cell; central memory CD4-positive, alpha-beta T cell; effector memory CD4-positive, alpha-beta T cell; T follicular helper cell |
| 5 | CD8 T cells | naive thymus-derived CD8-positive, alpha-beta T cell; central memory CD8-positive, alpha-beta T cell; effector memory CD8-positive, alpha-beta T cell; effector memory CD8-positive, alpha-beta T cell, terminally differentiated |
| 6 | Dendritic cells | conventional dendritic cell; plasmacytoid dendritic cell; CD1c-positive myeloid den-dritic cell |
| 7 | NK cells | CD16-negative, CD56-bright natural killer cell, human; CD16-positive, CD56-dim nat-ural killer cell, human; natural killer cell |
| 8 | Vascular endothelium | endothelial cell of artery; endothelial cell of venule; capillary endothelial cell; vein en-dothelial cell |
| 9 | Organ endothelium | endothelial cell of lymphatic vessel; vein endothelial cell of respiratory system; blood vessel endothelial cell |
| 10 | Smooth muscle | vascular associated smooth muscle cell; blood vessel smooth muscle cell; enteric smooth muscle cell; smooth muscle cell |
| 11 | Fibroblasts | myofibroblast cell; skin fibroblast; alveolar type 1 fibroblast cell; fibroblast |
| 12 | Macrophages | alveolar macrophage; elicited macrophage; macrophage |
| 13 | Lung epithelium | pulmonary alveolar type 1 cell; pulmonary alveolar type 2 cell; club cell; ciliated cell |
| 14 | Basal cells | basal cell; respiratory basal cell |
| 15 | Goblet cells | goblet cell; nasal mucosa goblet cell; intestine goblet cell |
| 16 | Intestinal epithelium | enterocyte; colonocyte; gut absorptive cell; intestinal epithelial cell |
| 17 | Nervous system | neuron; astrocyte; oligodendrocyte; microglial cell |
| 18 | Neural progenitors | neuroblast (sensu Vertebrata); oligodendrocyte precursor cell; Schwann cell |
| 19 | Stem/progenitor cells | hematopoietic multipotent progenitor cell; common myeloid progenitor; mesenchy-mal stem cell; stem cell |
| 20 | Regulatory T cells | regulatory T cell; CD4-positive, CD25-positive, alpha-beta regulatory T cell |
| 21 | ILC | innate lymphoid cell; immature innate lymphoid cell |
| 22 | Mural cells | pericyte; mural cell; perivascular cell |
| 23 | Blood lineage | erythrocyte; neutrophil; megakaryocyte; leukocyte |
| 24 | Connective tissue | chondrocyte; adipocyte; stromal cell |
| 25 | Secretory epithelium | mesothelial cell; choroid plexus epithelial cell; fallopian tube secretory epithelial cell; secretory cell |
| 26 | **tissue_general** | For example: brain, lung, blood |
| 27 | **sex** | male, female |
| 28 | **disease** | For example: lung adenocarcinoma, COVID-19 |
| 29 | **high_level_stage_name** | For example: embryonic stage, 80 years or older stage |

## Footnotes

1 ^1^https://github.com/karpathy/autoresearch

